# Gene Expression Network Analysis Provides Potential Targets Against SARS-CoV-2

**DOI:** 10.1101/2020.07.06.182634

**Authors:** Ana I. Hernández Cordero, Xuan Li, Chen Xi Yang, Stephen Milne, Yohan Bossé, Philippe Joubert, Wim Timens, Maarten van den Berge, David Nickle, Ke Hao, Don D. Sin

## Abstract

**BACKGROUND:** Cell entry of SARS-CoV-2, the novel coronavirus causing COVID-19, is facilitated by host cell angiotensin-converting enzyme 2 (ACE2) and transmembrane serine protease 2 (TMPRSS2). We aimed to identify and characterize genes that are co-expressed with *ACE2* and *TMPRSS2*, and to further explore their biological functions and potential as druggable targets.

**METHODS:** Using the gene expression profiles of 1,038 lung tissue samples, we performed a weighted gene correlation network analysis (WGCNA) to identify modules of co-expressed genes. We explored the biology of co-expressed genes using bioinformatics databases, and identified known drug-gene interactions.

**RESULTS:** *ACE2* was in a module of 681 co-expressed genes; 12 genes with moderate-high correlation with *ACE2* (r>0.3, FDR<0.05) had known interactions with existing drug compounds. *TMPRSS2* was in a module of 1,086 co-expressed genes; 15 of these genes were enriched in the gene ontology biologic process ‘Entry into host cell’, and 53 *TMPRSS2-*correlated genes had known interactions with drug compounds.

**CONCLUSION:** Dozens of genes are co-expressed with *ACE2* and *TMPRSS2*, many of which have plausible links to COVID-19 pathophysiology. Many of the co-expressed genes are potentially targetable with existing drugs, which may help to fast-track the development of COVID-19 therapeutics.

## INTRODUCTION

Since the outbreak of severe acute respiratory syndrome coronavirus (SARS coronavirus) in 2002 and 2003, coronaviruses have been considered to be highly pathogenic for humans ^1–3^. A new coronavirus (SARS-CoV-2) that spreads through respiratory droplets emerged in December of 2019 in Wuhan, Hubei province, China ^4^. Its rapid spread across China and the rest of the world ultimately resulted in the global COVID-19 pandemic, which was officially declared in March 2020 by the World Health Organization (WHO).

It has since been shown that SARS-CoV-2 shares a common host cell entry mechanism with the 2002-2003 SARS coronavirus ^5,6^. The viral genome encodes for multiple viral components, including the spike protein (S), which facilitates viral entry into the host cell ^7,8^. The S protein associates with the angiotensin-converting enzyme 2 (ACE2) to mediate infection of the target cells^9^. ACE2 is a type 1 transmembrane metallocarboxypeptidase that is an important negative regulator of the renin–angiotensin system (RAS). Once SARS-CoV-2 gains entry into epithelial cells, surface expression of ACE2 is rapidly downregulated in the infected cell^10^, which leads to an imbalance in angiotensin II-mediated signalling and predisposes the host to acute lung injury. SARS-CoV-2 also employs transmembrane serine protease 2 (TMPRSS2) to proteolytically activate the S protein, which is essential for viral entry into target cells^11^.

In view of the rapid spread and the mortality of COVID-19 worldwide, there is an urgent need to find effective treatments against this infection, especially for severe cases. However, the development of a vaccine and novel treatments may take months to years, requiring billions of dollars in investment and with no certainty of their ultimate success. Bioinformatic approaches, however, can rapidly identify relevant gene-drug interactions that may contribute to the understanding of the mechanisms of viral infection and reduce the time to finding potential drug targets and existing drugs that could be repurposed for this indication. Here, we performed a gene expression network analysis on data generated in the Lung eQTL Consortium Cohort to investigate the mechanisms of *ACE2* and *TMPRSS2* expression in lung tissue. We identified potential targets to be explored as possible treatments for COVID-19. We hypothesized that the mechanisms associated with *ACE2* and *TMPRSS2* likely encompass protein coding genes involved in the pathogenesis of COVID-19.

## RESULTS

The Lung eQTL Consortium cohort used in this gene network analysis is described in **Table 1. Supplementary Fig. S1** shows the expression levels of *ACE2* and *TMPRSS2* in the three centres that are part of the Lung eQTL Consortium (see methods); *ACE2* had low to moderate expression levels in lung tissue; whereas *TMPRSS2* was highly expressed. Based on the study cohort lung expression profile, we determined that *ACE2* and *TMPRSS2* were contained in distinct modules. The module containing *ACE2* (*ACE2* module) included 681 unique genes, while the modules containing *TMPRSS2* (*TMPRSS2* module) encompassed 1,086 unique genes (**Supplementary Tables S1 and S2**). Only 41 genes were found in both modules. The hub gene for the *ACE2* module was *TMEM33*, and hub gene for the *TMPRSS2* module was *PDZD2* (see methods for the definition of ‘hub gene’). **Fig. 1** shows the top 50 genes with the highest connectivity to *ACE2* and *TMPRSS2* within their respective modules, based on the weighted gene correlation network analysis (WGCNA) analysis.

**Table 1.**
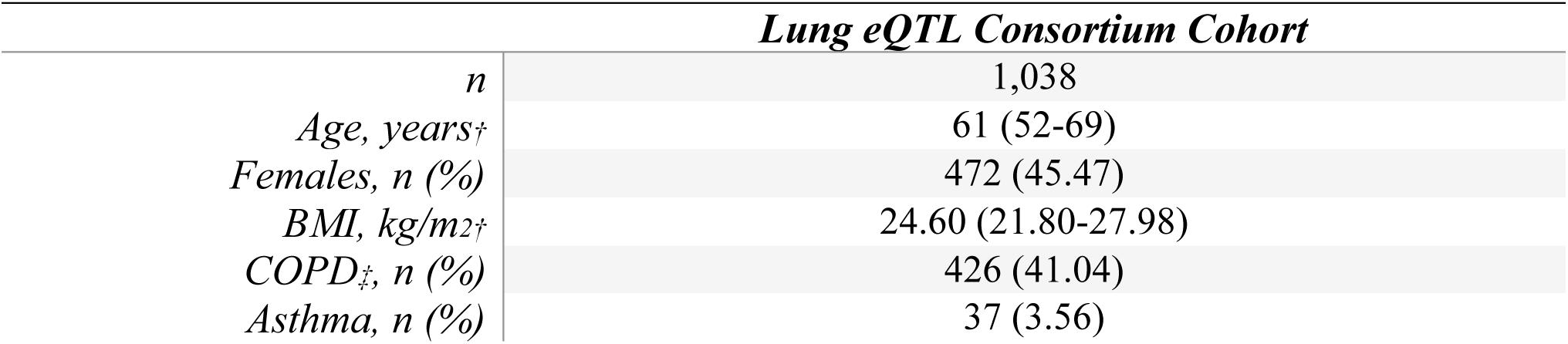
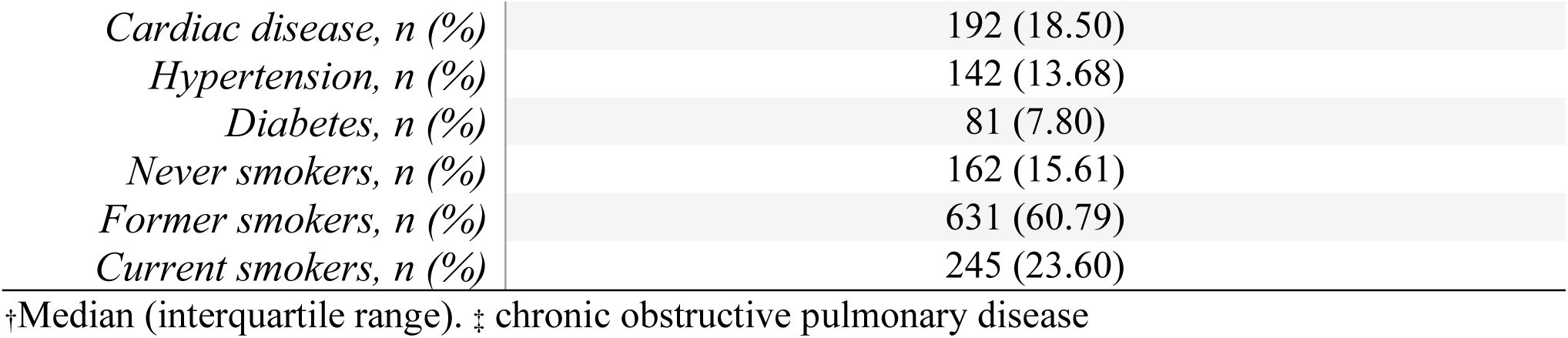
Study cohort demographics.

| *Lung eQTL Consortium Cohort* |  |
| --- | --- |
| *n* | 1,038 |
| *Age, years*<sup>†</sup> | 61 (52-69) |
| *Females, n (%)* | 472 (45.47) |
| *BMI, kg/m<sup>2</sup>*<sup>†</sup> | 24.60 (21.80-27.98) |
| *COPD*<sup>‡</sup>, *n (%)* | 426 (41.04) |
| *Asthma, n (%)* | 37 (3.56) |
| *Cardiac disease, n (%)* | 192 (18.50) |
| *Hypertension, n (%)* | 142 (13.68) |
| *Diabetes, n (%)* | 81 (7.80) |
| *Never smokers, n (%)* | 162 (15.61) |
| *Former smokers, n (%)* | 631 (60.79) |
| *Current smokers, n (%)* | 245 (23.60) |

†Median (interquartile range). ‡ chronic obstructive pulmonary disease

**Figure 1.**
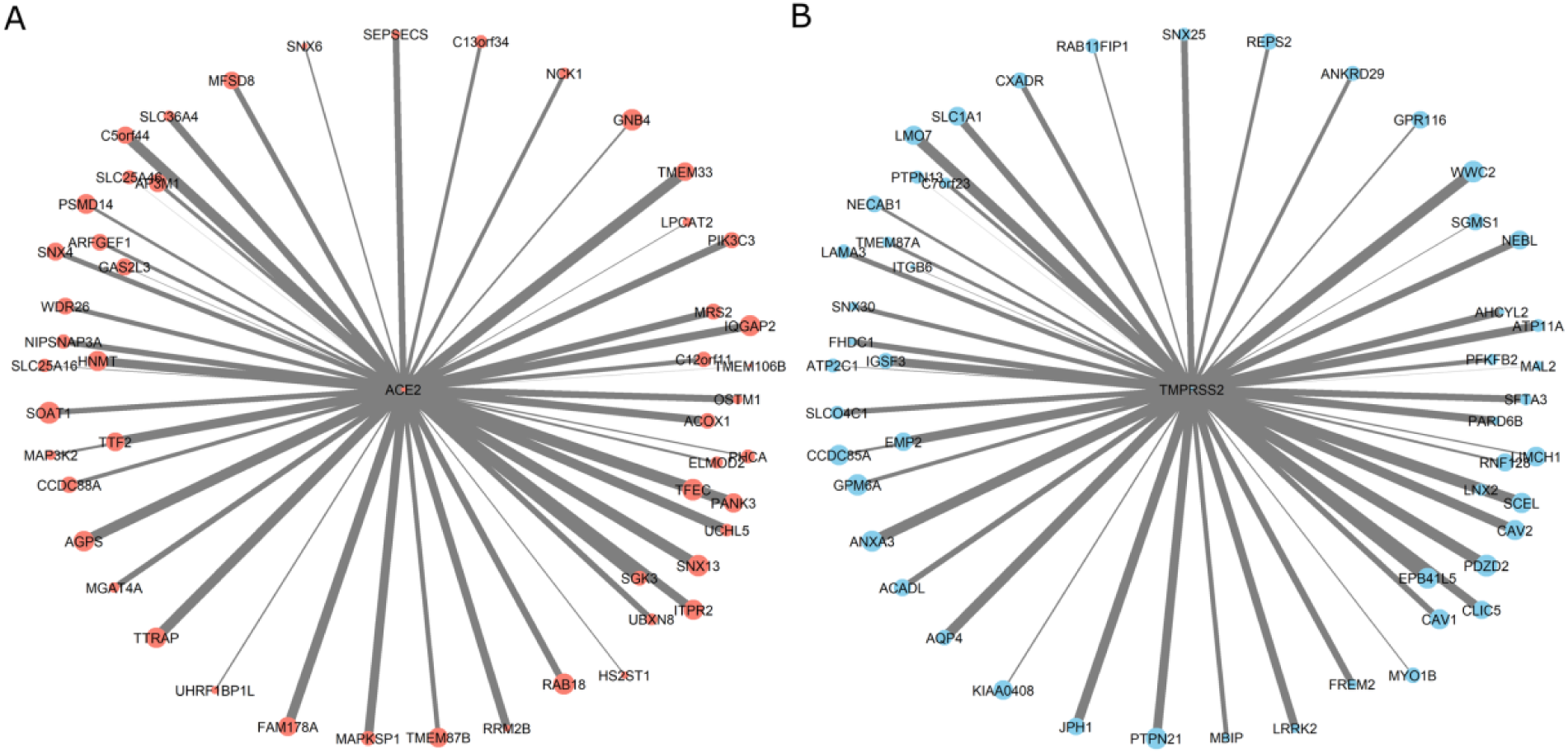
*ACE2* and *TMPRSS2* expression modules. The center of each graph represents *ACE2* (A) or *TMPRSS2* (B), the circles at the edges represent the top 50 genes with the highest connectivity to *ACE2* or *TMPRSS2* based on the WGCNA analysis. The circle size represents the size of each gene node in their respective modules. The arm thickness represents the relative strength of the connection to the *ACE2* or *TMPRSS2* expression.

### ACE2 module

The median module membership (MM) (see methods) across the genes in the *ACE2* module was 0.40, and the minimum and maximum values were 0.002 and 0.79, respectively. The MM for *ACE2* was 0.25. We utilized genes in the *ACE2* module to execute a pathway enrichment analysis, which showed significant enrichment of four Kyoto Encyclopedia of Genes and Genomes (KEGG) pathways (Lysosome, Metabolic pathways, N-Glycan biosynthesis and Endocytosis) (**Supplementary Table S3**) and 34 gene ontology (GO) biologic processes (*FDR<0*.*05*) (**Supplementary Table S4**); however *ACE2* was not part of the enriched pathways or processes.

### ACE2-correlated genes

The expression of 646 genes in the *ACE2* module was significantly correlated with *ACE2* levels (*FDR<0*.*05*), and only two of those genes were negatively correlated with *ACE2*. The range of their correlation coefficient (r) with *ACE2* expression level in lung tissue is shown in **Supplementary Fig. S2**. Although a large proportion of genes were significantly related to *ACE2* expression levels, only 76 genes had moderate or high correlations (r>0.3).

The *PCCB* gene was most strongly correlated with *ACE2* expression (r=0.45, **Supplementary Table S1**). Of the top 10 genes most strongly correlated with *ACE2*, three genes (*PCCB, PIGN* and *ADK*) were part of the KEGG ‘metabolic pathway’ which showed enrichment with *ACE2* module genes (**Supplementary Table S3**). Furthermore, out of the top 10 genes, four genes (*ITPR2, LONP2, ADK* and *WDFY3*) were found in multiple GO processes that were enriched with *ACE2* module genes (**Supplementary Table S4**).

We identified 76 genes that showed moderate correlation (r>0.3) with *ACE2* expression. Of these, 48 genes had biological and/or druggability information available (details are presented in **Supplementary Table S1**). We used these genes to construct a ‘map’ of biological information (**Supplementary Fig. S3)**. Based on the druggability scores, we identified 13 genes (*GART, DPP4, PIGF, HDAC8, MDM2, SOAT1, IDE, BCAT1, SLC11A2, ADK, KLHL8, IL13RA2* and *ITPR2*) that are known drug targets or are part of a key pathway that is targeted by a drug compound (see methods for details on druggability scores). Out of the13 genes with druggability scores, 12 were found to have of known drug-gene interactions (**Table 2**).

**Table 2.** Drug-gene interactions of *ACE2*-correlated genes.

| *Gene* | **Druggability score<sup>†</sup>** | **No. of known drug-gene interactions<sup>‡</sup>** | **r (ACE2)** | ***p*** | ***FDR*** |
| --- | --- | --- | --- | --- | --- |
| *ITPR2* | Tier 3 | 5 | 0.42 | $1.88 \times 10^{-46}$ | $8.62 \times 10^{-44}$ |
| *ADK* | Tier 2 | 12 | 0.37 | $9.75 \times 10^{-36}$ | $9.91 \times 10^{-34}$ |
| *GART* | Tier 1 | 2 | 0.36 | $7.77 \times 10^{-34}$ | $4.74 \times 10^{-32}$ |
| *SOAT1* | Tier 2 | 8 | 0.35 | $1.47 \times 10^{-32}$ | $6.71 \times 10^{-31}$ |
| *SLC11A2* | Tier 2 | 1 | 0.34 | $1.28 \times 10^{-30}$ | $4.52 \times 10^{-29}$ |
| *IDE* | Tier 2 | 3 | 0.34 | $8.11 \times 10^{-30}$ | $2.06 \times 10^{-28}$ |
| *BCAT1* | Tier 2 | 6 | 0.33 | $7.58 \times 10^{-29}$ | $1.61 \times 10^{-27}$ |
| *DPP4* | Tier 1 | 48 | 0.32 | $5.95 \times 10^{-27}$ | $9.90 \times 10^{-26}$ |
| *MDM2* | Tier 1 | 2 | 0.32 | $9.56 \times 10^{-27}$ | $1.56 \times 10^{-25}$ |
| *PIGF* | Tier 1 | 1 | 0.32 | $9.87 \times 10^{-27}$ | $1.56 \times 10^{-25}$ |
| *HDAC8* | Tier 1 | 27 | 0.32 | $1.21 \times 10^{-26}$ | $1.87 \times 10^{-25}$ |
| *IL13RA2* | Tier 3 | 1 | 0.32 | $1.30 \times 10^{-26}$ | $1.98 \times 10^{-25}$ |

<sup>†</sup>from Finan et al <sup>12</sup> <sup>‡</sup>from Drug-Gene Interaction Database (DGIdb) <sup>13</sup>. r(ACE2): Pearson correlation coefficient between gene and ACE2 expression.

### TMPRSS2 module

*TMPRSS2* demonstrated a MM of 0.27 (**Supplementary Table S2**). Genes in the *TMPRSS2* module were enriched in multiple KEGG pathways (**Supplementary Table S5**) and GO biologic processes (**Supplementary Table S6**). Five of the GO biologic processes identified in this study, including ‘entry into host cell’, also contained *TMPRSS2* (**Table 3**).

**Table 3.** GO biological processes involving *TMPRSS2* and enriched in the *TMPRSS2* module.

| **GO biological process** | **Overlap<sup>†</sup>** | **Enrichment Ratio** | ***p*** | ***FDR*** |
| --- | --- | --- | --- | --- |
| Endocytosis | 68/675 | 1.98 | $5.39 \times 10^{-08}$ | $5.38 \times 10^{-06}$ |
| Vesicle-mediated transport | 149/1,942 | 1.51 | $1.33 \times 10^{-07}$ | $1.17 \times 10^{-05}$ |
| Import into cell | 74/792 | 1.83 | $2.88 \times 10^{-07}$ | $2.22 \times 10^{-05}$ |
| Receptor-mediated endocytosis | 34/287 | 2.32 | $4.13 \times 10^{-06}$ | $2.26 \times 10^{-04}$ |
| Entry into host cell | 15/134 | 2.20 | $3.42 \times 10^{-03}$ | $4.59 \times 10^{-02}$ |

<sup>†</sup>Number of genes identified by our research over the total number of genes in the GO biological process

### TMPRSS2-correlated genes

We found that 864 unique genes in the *TMPRSS2* module were positively correlated with the *TMPRSS2* expression level in lung tissue (*FDR<0*.*05*), while 73 demonstrated a negative relationship with the gene. The absolute r ranged from 0.06 to 0.72, with *FHDC1* expression showing the strongest correlation with *TMPRSS2* (r=0.72) (**Supplementary Table S2**). Next, we identified 368 genes that were moderately or highly correlated with *TMPRSS2* gene expression levels (r>0.30), of those 78 were drug targets or were part of key pathways that could be targeted by drug compounds (see methods). The genes are shown in **Fig. 2**, grouped based on the availability of biological information. The A4 group contained the genes with the most amount of biological information in the explored bioinformatics databases. Most genes in **Fig. 2** only had information on human phenotypes (A1 group); details on the genes biological information are presented in the **Supplementary Table S2**.

**Figure 2.**
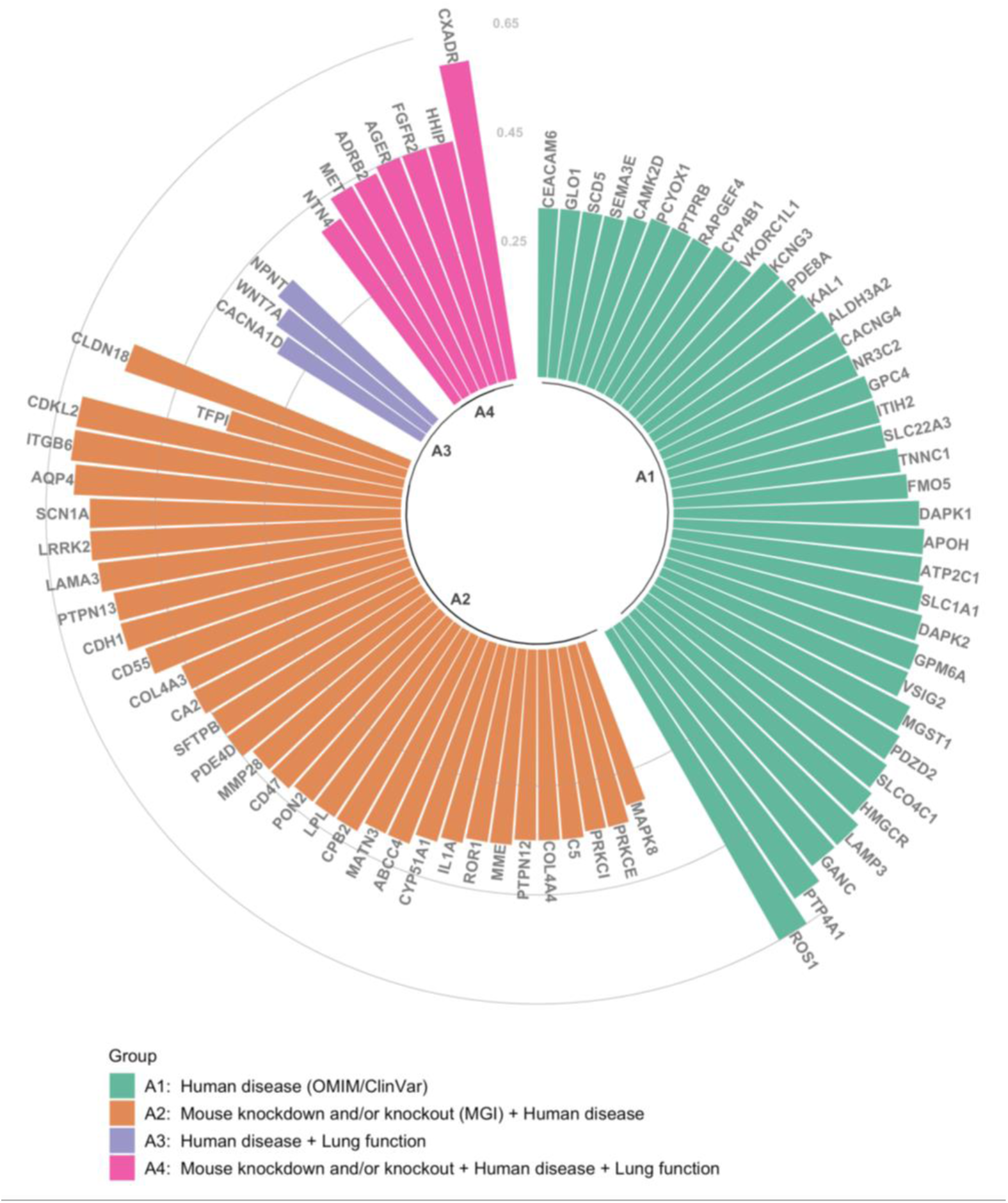
Correlation level and annotation of *TMPRSS2*-correlated genes. Each bar represents a single gene (all with druggability scores Tier 1-3^12^), and Pearson correlation coefficient (r) between the gene and *TMPRSS2* within the module is shown on the y axis. Colours of bars represent combined biological information: green (group A1) represents genes related to human diseases based Online Mendelian Inheritance in Man (OMIM) and ClinVar databases; orange (group A2) are genes associated with human diseases, which also have phenotypic information on knockdown or knockout mouse models based on Mouse Genome Informatics (MGI) database; purple (group A3) represents genes associated with human diseases and with genetic variants associated to lung function traits^14^; pink (group A4) represents genes associated with a human disease, with phenotypic information on knockdown or knockout mouse, and genetic variants associated with lung function.

We later explored the drug-gene interactions of the genes described in **Fig. 2**; 53 of these genes were found to interact with known drugs. Furthermore, 21 genes with gene-drug interactions (**Table 4**) were enriched in the GO biological processes that related to *TMPRSS2* (**Table 3**). The **Table 4** includes four out of the 15 genes that are part of the ‘*Entry into host cell*’ biological process (*CD55, CDH1, ITGB6* and *MET*).

**Table 4.**
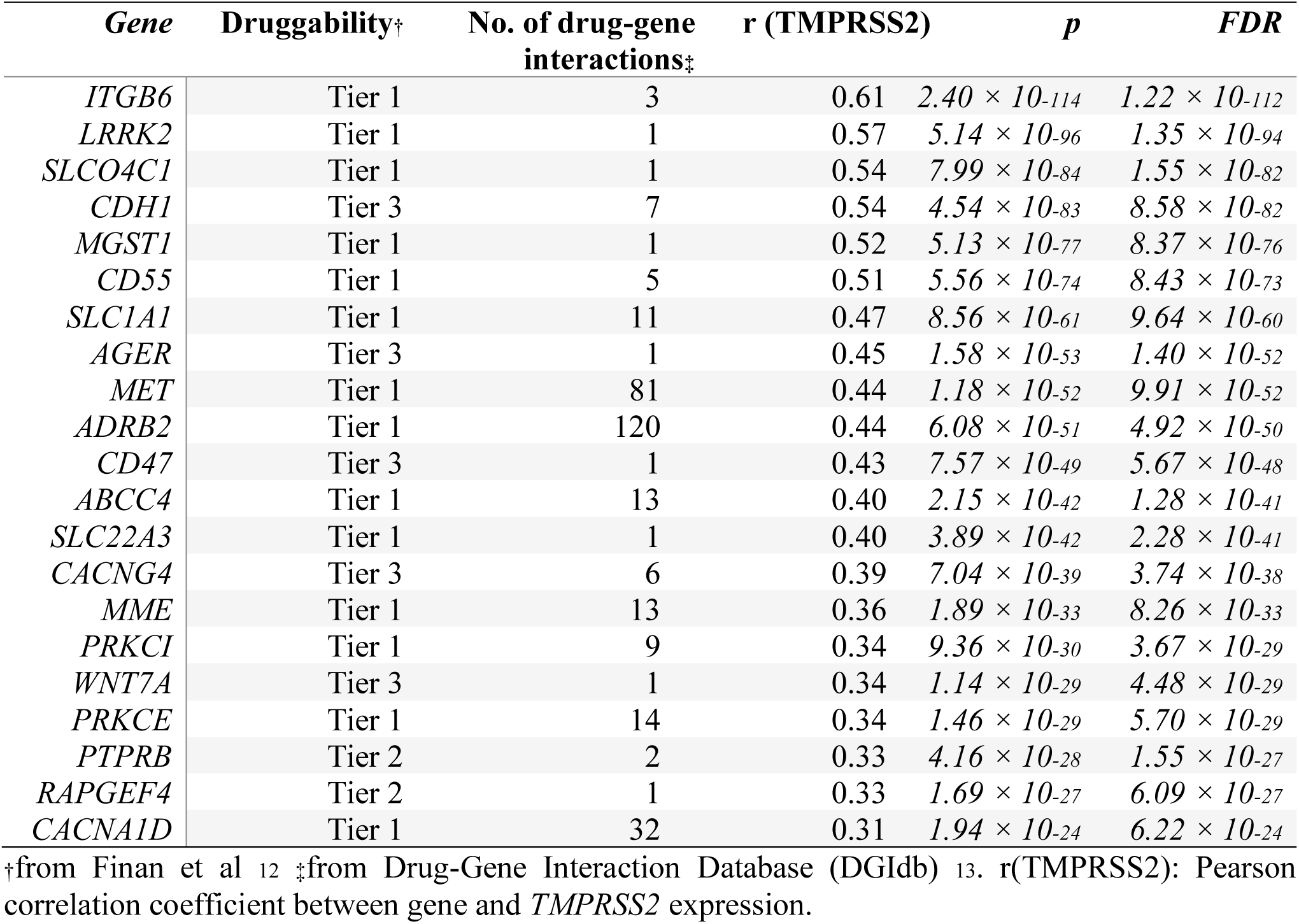
Drug-gene interactions of *TMPRSS2*-correlated genes.

### Differential expression of ACE2- and TMPRSS2-correlated genes

We investigated the effects of risk factors for COVID-19 on the expression of the genes shown in **Table 2** and **Table 4**. The full list of differential expressed genes (*FDR<0*.*05*) with known drug-gene interactions is presented in **Supplementary Table S7**. Some illustrative examples are shown in **Fig. 3**, including the effect of chronic obstructive pulmonary disease (COPD) on *IL13RA2* expression (**Fig. 3A**), and the effect of smoking on *ADK, DPP4, MGST1, CD55* and *ITGB6* expression (**Fig. 3B-F**).

**Figure 3.**
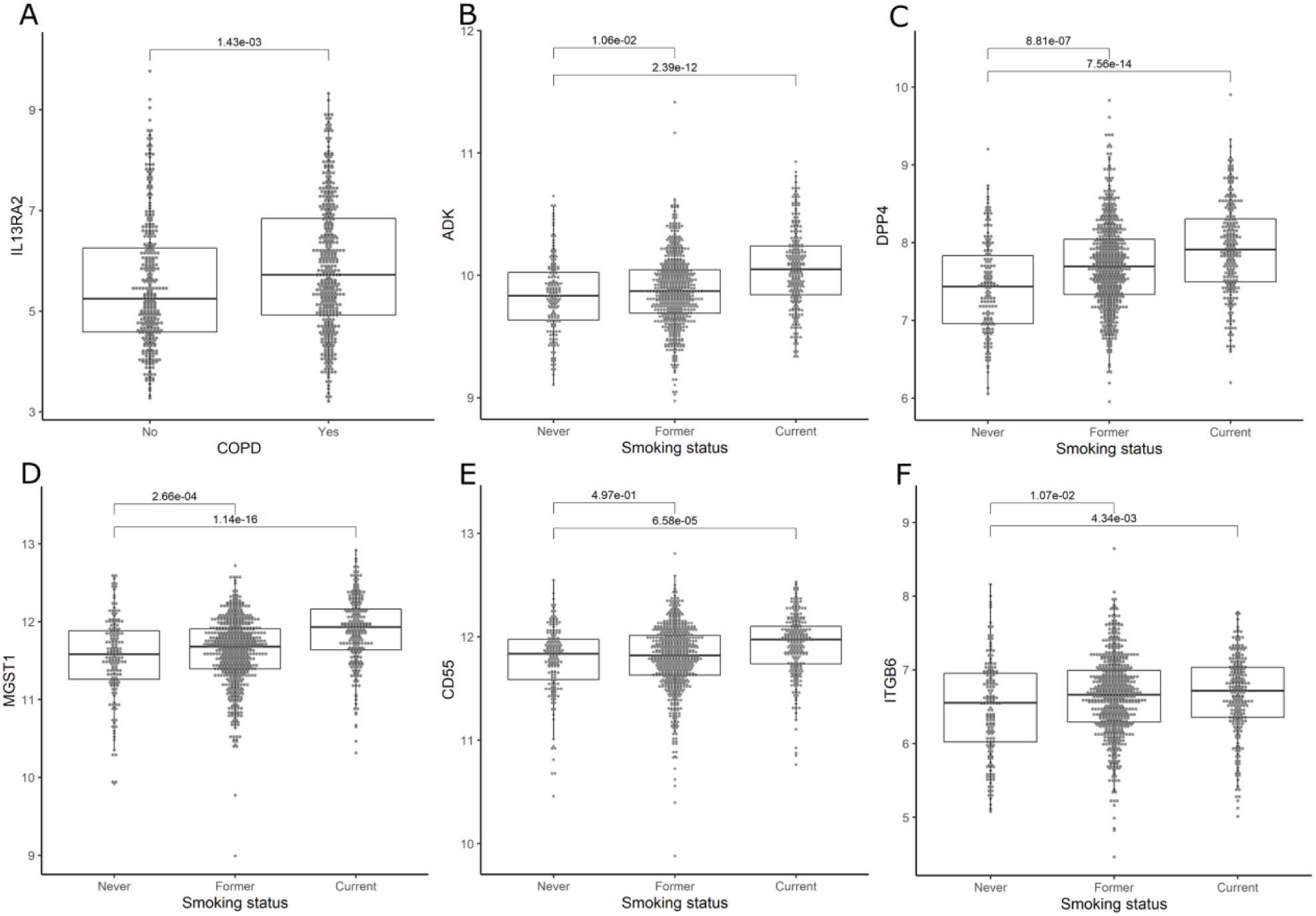
Effects of COVID-19 risk factors on lung tissue gene expression. y axes represent the expression level in log2(counts per million) in lung tissue for *ACE2*-correlated genes (*IL13RA2* [A], *ADK* [B], *DPP4* [C]) and *TMPRSS2*-correlated genes (*MGST1* [D], *CD55* [E], *ITGB6* [F]). Boxes are median expression ± interquartile range respectively. Numbers at the top of each box plot are FDR obtained from the robust linear regressions.

## DISCUSSION

There is a scarcity of therapeutic treatments specific for this virus and for severe COVID-19 pneumonia. *ACE2* and *TMPRSS2* are key proteins involved in the cellular entry mechanism of SARS-Cov-2 to infect the lungs of the human host. Because one of the rate-limiting step in this process is the overall availability of these proteins on surface of lung epithelial cells 15, careful evaluation of *ACE2* and *TMPRSS2* biology may enable identification of possible therapeutic targets against SARS-Cov-2 infection. In this study, by using a network analysis of genome-wide gene expression in lung tissue, we were able to identify a set of genes that may interact with *ACE2* and *TMPRSS2*, and thus may be drug targets.

One notable gene was *ADK*. This gene is a key regulator of extracellular and intracellular adenine nucleotides ^16,17^. ADK inhibition attenuates lung injury in mice ^18^, while in humans, cigarette exposure upregulates expression of *ADK* in lung tissue. We speculate a role for *ADK* in COVID-19, postulating that increased ADK may increase adenosine concentration in the lungs which in turn can enhance viral replication. Previous work has shown that silencing *ADK* decreased influenza replication in an *in vitro* model^19^. Another study showed that ADK can activate didanosine^20^, a dideoxynucleoside analogue of adenosine that inhibits retro-transcription and is used in the treatment of HIV. Although this drug was recently nominated for drug repurposing as a potential treatment against COVID-19^21^, the biology of this drug is complex, particularly given the detrimental effect of ADK on lung injury.

Another *ACE2*-correlated gene that emerged from this study was *DPP4. DPP4* encodes the dipeptidyl-peptidase 4 (DPP-4) glycoprotein, which plays a major role in glucose and insulin metabolism and is linked to diabetes, now established as a key risk factor for severe COVID-19 including mortality^22^. DPP-4 is the functional receptor for the Middle East Respiratory Syndrome (MERS) coronavirus and interacts with dozens of drugs. DPP-4 inhibitors, which are used in the treatment of diabetes, appear to reduce macrophage infiltration and insulin resistance but have not been shown to increase the risk of infection in diabetic patients^23^. However, the effects of DPP-4 inhibitors on the immune response are not well understood. Because of the similarities between MERS and SARS-Cov-2, this is an interesting potential target, particularly for patients with diabetes.

Another interesting target is *IL13RA2*, which encodes the alpha-2 receptor subunit for interleukin-13 (IL-13). The IL-13 pathway has immunoregulatory functions and is implicated in asthma, idiopathic pulmonary fibrosis (IPF)^24^ and COPD ^25,26^. The IL-13 pathway can activate Janus kinase 2 (JAK2) while the inhibition of JAK2 blocks SARS-CoV-2 viral entry^27^. *IL13RA2* interacts with cintredekin besudotox, a drug compound that is formed by cross-linking IL-13 with *Pseudomonas* exotoxin-A and induces apoptosis by targeting cells that express the IL-13 receptor. Both the IL-13 and DPP-4 pathways could be intriguing possibilities for novel COVID-19 therapeutics.

The *HDAC8* gene is an exciting potential target because of its role in pulmonary fibrosis (PF) and its interaction with histone deacetylase (HDAC) inhibitors. HDAC inhibitors have shown promise against fibrotic diseases^28^. The overexpression of HDACs is suggested to contribute to the process of bronchiolization in patients with IPF^29^ Viral infection increases the risk of PF^30^ and it is reported that *HDAC8* inhibition ameliorates PF ^31^; moreover we found that cigarette exposure, a known risk factor for both COVID-19 and IPF, increases the expression of *HDAC8* in lung tissue. Therefore, targeting the PF mechanisms through HDAC inhibitors pose an interesting therapy to further explore.

The ‘entry into host cell’ biological process was enriched with genes from the *TMPRSS2* module. Furthermore, the *CD55* or complement decay-accelerating factor, an inhibitor of complement activation, is one of the few genes that was part of this process. The complement system has a major role in the immune response to viruses and triggers a proinflammatory cascade ^32^. CD55, which is highly expressed in lung tissue, prevents the formation of C3 convertase ^33^ and therefore also inhibits the formation of C3 complement. C3-deficient mice show less respiratory dysfunction and lower levels of cytokines and chemokines in lungs in response to SARS-CoV^34^. Thus, it is possible that preventing the formation of C3 via CD55 could be beneficial in COVID-19. Fortunately, known compounds such as chloramphenicol already exist that specifically target CD55 ^31,35^.

As noted above, we have identified a set of genes that interact with potential therapeutic targets, which could be explored as treatments against COVID-19. The main strength of our study is the large number of lung tissue specimens with detailed clinical phenotypic data. This allowed us to not only identify genes related to *ACE2* and *TMPRSS2* expression, but also to determine the effects of various clinical factors on the lung tissue expression of these genes. However, there were limitations to this study. First, we used an *in-silico* approach to identify *ACE2* and *TMPRSS2* correlated genes, but we did not confirm these association *in vivo* nor determine how these correlated genes physically interacted with *ACE2* and *TMPRSS2*. Second, we identified the most promising drugs based on drug-gene interactions from bioinformatic databases, but we are yet to test their effects on gene and/or protein expression in *in vitro* experiments. Third, the lungs of our study cohort were not exposed to SARS-CoV-2, therefore it is possible that the gene expression of these key identified genes in lung tissue could be changed upon SARS-CoV-2 infection. Lastly, the cohort used for gene expression was of European ancestry and the results may not be generalizable to other ethnic groups, which is of critical importance in a global pandemic.

In summary, *ACE2* and *TMPRSS2* gene networks contained genes that could contribute to the pathophysiology of COVID-19. These findings show that computational *in silico* approaches can lead to the rapid identification of potential drugs, which could be repurposed as treatments against COVID-19. Given the exponential spread of COVID-19 across the globe and the unprecedented rise in deaths, such rapidity is necessary in our ongoing fight against the pandemic.

## METHODS

### Lung expression Quantitative Trait Loci (eQTL) Consortium Cohort and gene expression

Using microarray, gene expression profiles of 43,466 non-control probe sets (GEO platform GPL10379) were obtained from lung tissue samples in the Lung eQTL Consortium Cohort. Briefly, samples from this cohort included whole non-tumour lung tissue samples from 1,038 participants of European ancestry who underwent surgical lung resection. Tissue samples were collected based on the Institutional Review Board guidelines at three different institutions: The University of British Columbia (UBC), Laval University and University of Groningen. This study was approved by the ethics committees within each institution. A full description of the cohort and quality controls is provided by Hao and colleagues 36.

### Gene expression network analysis

Using the WGCNA^37^ R package, we explored gene networks correlated to *ACE2* and *TMPRSS2* in order to identify potential interactions in the Lung eQTL Consortium cohort. WGCNA clusters co-expressed genes into networks and creates “modules”, which are defined as groups of highly interconnected genes. For this analysis we identified signed consensus modules among the three centres in our study cohort. Briefly, WGCNA generated a signed co-expression matrix based on the correlation between genes, which later was transformed into an adjacent matrix by raising the co-expression to a soft threshold power (β). For our study we used a β=6 and a minimum module size of 100 probe sets. A consensus network was built by identifying the overlap of all input datasets. For each probe set in the modules a ‘Module Membership’ (MM) was calculated by correlating the gene’s expression with the respective module’s expression (eigengene), i.e. the first principal component of each module gene expression profile; the gene with the highest MM was termed the ‘hub gene’.

### Enrichment analysis and correlations of ACE2 and TMPRSS2 modules

Enrichment analysis of KEGG pathways and GO biological processes was performed using the genes in the modules containing *ACE2* (*ACE2* module) and *TMPRSS2* (*TMPRSS2* module). Significant enrichment was established at *FDR<0*.*05*. For each gene in the *ACE2* and *TMPRSS2* modules, we determined the Pearson correlation between the expression level of the gene and that of *ACE2* or *TMPRSS2*. We calculated correlation coefficients for the three centres separately and then combined them using correlation meta-analysis via the R package metafor ^38^. Significant correlations were set at *FDR<0*.*05* and in the downstream analyses, we focused on genes that correlated to *ACE2* or *TMPRSS2* with r>0.30.

### Drug-gene interactions and biological information of ACE2 and TMPRSS2 correlated genes

We cross-referenced the *ACE2* and *TMPRSS2* correlated genes with the Mouse Genome Informatics (MGI), the Online Mendelian Inheritance in Man (OMIM), and the ClinVar databases in order to identify biologically relevant genes. We determined druggability scores according to methods of Finan et al^12^. Tier 1 refers to genes that are targets of small molecules and/or biotherapeutic drugs; Tier 2 score indicates gene encoding targets with a known bioactive drug-like small molecule binding partner and ≥50% identity (over ≥75% of the sequence) with an approved drug target; and Tier 3 denotes protein coding genes with similarities to drug targets and are members of key druggable gene families. We also interrogated the Drug-Gene Interaction database (DGIdb)^13^ of the genes. DGIdb defines drug-gene interaction as a known interaction (i.e.: inhibition, activation) between a known drug compound and a target gene.

### Differential expression of ACE2 and TMPRSS2 correlated genes

We investigated the effects of possible risk factors for COVID-19 severity (e.g. smoking, diabetes, asthma, COPD, cardiac disease, and hypertension) on the expression of druggable genes that were correlated with *ACE2* or *TMPRSS2*. We first combined the gene expression from the three centres using ComBat from the R package sva to correct for any batch effect^39^ Then, the differential expression was assessed for each gene-risk factor pair by a robust linear regression using the package MASS^40^ in R, in which the dependent variable was the gene expression and the explanatory variable was the risk factor. The differential expression analysis on smoking was adjusted for sex and age, and the analyses on diabetes, COPD and cardiac disease and hypertension were additionally adjusted for smoking status. We set statistically significant differential expression *FDR<0*.*05*.

## DATA AVAILABILITY

The full results obtained in this analysis are provided in the Supplementary Tables associated to this manuscript.

## Acknowledgments

The eQTL data from Laval University were generated from tissues obtained through the Quebec Research Respiratory Network Biobank, IUCPQ site. We acknowledge Compute Canada and WestGrid, which provided computational resources to conduct this research.

## Authors’ contributions

A.I.H.C. wrote the draft of the manuscript. X.L. conducted the main analyses with the impute of A.I.H.C and revised the manuscript. C.X.Y. and S.M. revised the manuscript. Y.B., P.J., W.T., M.B., D.N., and K.H. provided lung expression data and revised the manuscript. D.D.S. supervised this study and revised the manuscript.

## ADDITIONAL INFORMATION

### Competing interests

S.M. reports personal fees from Novartis and Boehringer-Ingelheim, outside the submitted work. W.T. reports fees to Institution from Roche-Ventana, AbbVie, Merck-Sharp-Dohme and Bristol-Myers-Squibb, outside the submitted work. M.B. reports research grants paid to University from Astra Zeneca, Novartis, outside the submitted work. D.D.S. reports research funding from AstraZeneca and received honoraria for speaking engagements from Boehringer Ingelheim and AstraZeneca over the past 36 months, outside of the submitted work.

### Funding

A.I.H.C. and S.M. are supported by MITACS Accelerate grant. D.D.S. holds the De Lazzari Family Chair at HLI and a Tier 1 Canada Research Chair in COPD. Y.B. holds a Canada Research Chair in Genomics of Heart and Lung Diseases

