## Supplementary Figures for "Gene Expression Network Analysis Provides Potential Targets Against SARS-CoV-2"

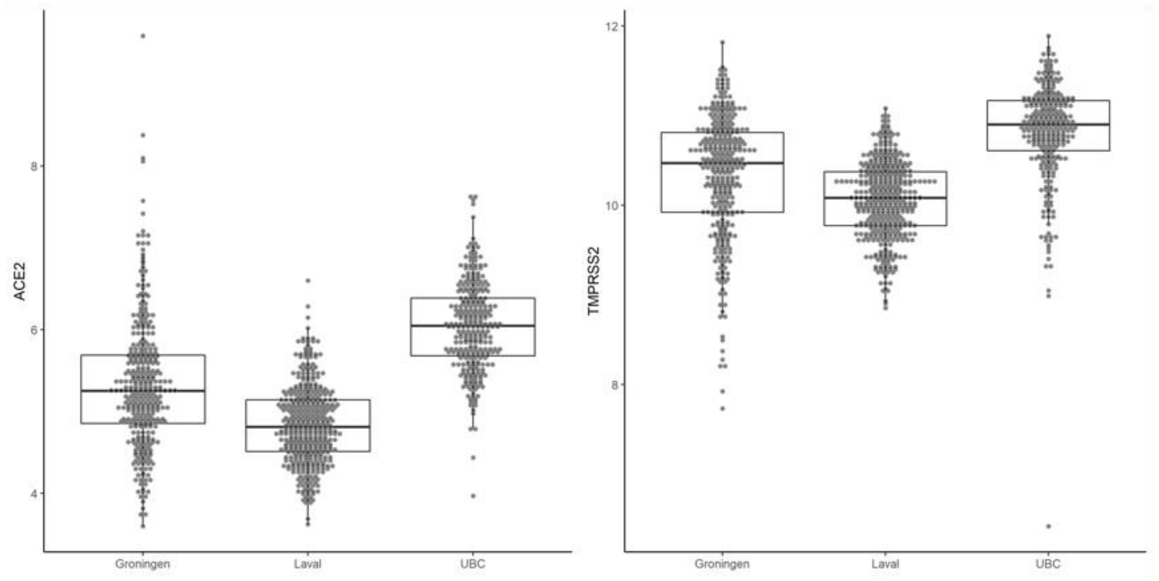


**Supplementary Figure S1.** ACE2 and TMPRSS2 expression level in lung tissue. The y-axis of the plots represents gene expression levels of ACE2 (A) and TMPRSS2 (B) and the x-axis represent each of the centres. The boxes and the horizontal line inside them, represent the interquartile range and median expression, respectively. The dots across the boxes represent the distribution of the gene expression.

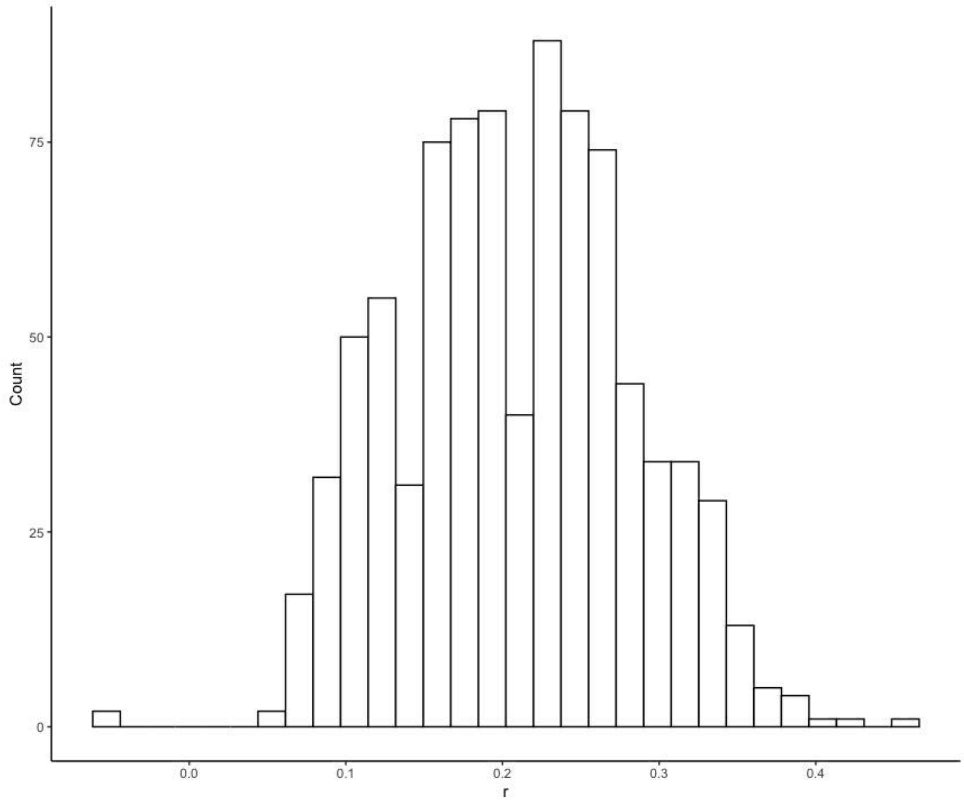


**Supplementary Figure S2.** Distribution of Pearson correlation coefficients (r) between lung tissue ACE2 expression and the 646 genes that are part of the ACE2 module (FDR<0.05). The y axis represents the counts of genes, and the x axis represents the correlation coefficients of genes with ACE2 expression.

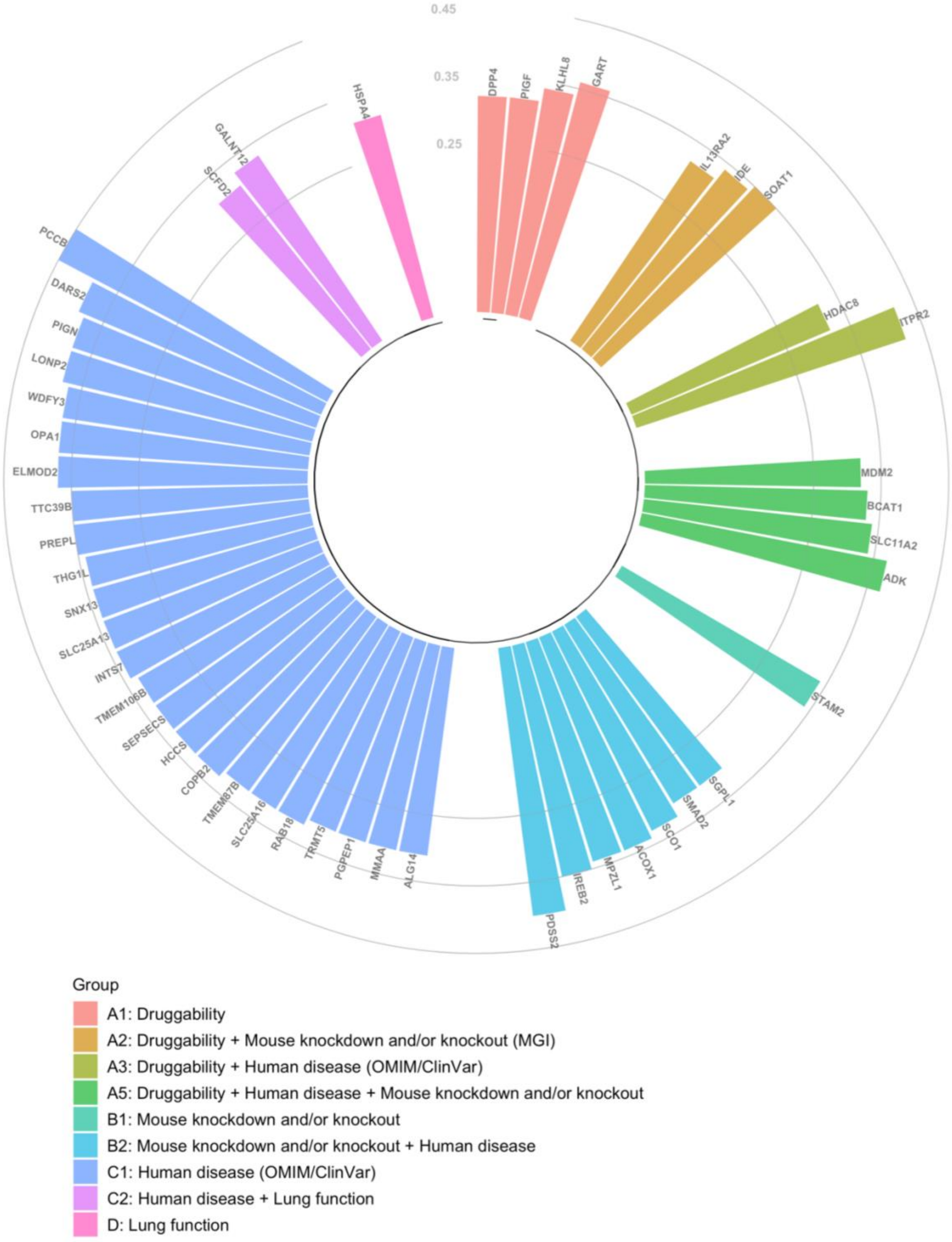


**Supplementary Figure S3.** Correlation level (y-axis) and annotation for ACE2 correlated genes. Each bar represents a single gene, and Pearson correlation coefficient (r) between the gene and TMPRSS2 within the module is shown on the y axis. Colours of bars represent combined biological information as described in the ‘Group’ information provided below the figure.
